# Accessing mitochondrial protein import in living cells by protein microinjection

**DOI:** 10.1101/2020.09.30.317412

**Authors:** Andrey Bogorodskiy, Ivan Okhrimenko, Ivan Maslov, Nina Maliar, Dmitry Burkatovskiy, Florian von Ameln, Alexey Schulga, Philipp Jakobs, Joachim Altschmied, Judith Haendeler, Alexandros Katranidis, Alexey Mishin, Valentin Gordeliy, Georg Büldt, Wolfgang Voos, Thomas Gensch, Valentin Borshchevskiy

## Abstract

Mitochondrial protein biogenesis relies almost exclusively on the expression of nuclear-encoded polypeptides. The current model postulates that most of these proteins have to be delivered to their final mitochondrial destination after their synthesis in the cytoplasm. However, the knowledge of this process remains limited due to the absence of proper experimental real-time approaches to study mitochondria in their native cellular environment. We developed a gentle microinjection procedure for fluorescent reporter proteins allowing a direct non-invasive study of protein transport in living cells. As a proof of principle, we visualized potential-dependent protein import into mitochondria inside intact cells in real-time. We validated that our approach does not distort mitochondrial morphology and preserves the endogenous expression system as well as mitochondrial protein translocation machinery. We observed that a release of nascent polypeptides chains from actively translating cellular ribosomes by puromycin strongly increased the import rate of the microinjected preprotein. This suggests that a substantial amount of mitochondrial translocase complexes were involved in co-translational protein import of endogenously expressed preproteins. Our protein microinjection method opens new possibilities to study the role of mitochondrial protein import in cell models of various pathological conditions as well as aging processes.

## Introduction

Mitochondria are among the most important organelles in eukaryotic cells producing the bulk of ATP, which fuels many vital biochemical and physiological processes in the cell. In mammalian cells, only 13 mitochondrial proteins are produced inside the mitochondria while the remaining vast majority of 1000-1500 proteins are expressed externally (Anderson et al., 1981; Pagliarini et al., 2008), translated at cytosolic ribosomes and later imported into the organelle. An extensive transport and sorting network is therefore required for mitochondrial protein biogenesis. At least five pathways described so far are responsible for sorting imported proteins into different mitochondrial compartments: outer/inner mitochondrial membrane (OMM/IMM), intermembrane space, and matrix. Four of them use the TOM (translocase of the outer mitochondrial membrane) complex as a gateway (Wiedemann and Pfanner, 2017). Protein import in the mitochondria requires the recognition of specific amino acid mitochondrial targeting sequences (MTS) of the precursor protein (preprotein) by components of the respective translocase complexes.

Experimental approaches up to date have yielded remarkable progress in defining the functional mechanism of mitochondrial protein import. Most experiments were done *in vitro* on the isolated mitochondria – essentially restricting the analyzed import process to a post-translational fashion outside of the genuine cellular environment. However, these experiments clearly established that mitochondrial protein import can principally function in a purely post-translational manner, and they do not necessarily need the cytosolic co-factors (Becker et al., 1992). Under saturating preprotein concentrations, rather high import rates can be achieved (Lim et al., 2001), showing that the post-translational import rates are high enough to avoid mitochondrial proteins accumulation in the cytoplasm.

Experiments performed in living cells have pointed to a large variety of cytosolic factors affecting the targeting of mitochondrial proteins in the post-translational import situation (Becker et al., 2019), in particular molecular chaperones Hsp70 and Hsp90 (Young et al., 2003). In addition, the chaperone co-factors from the Hsp40 protein family (Opaliński et al., 2018) are involved in the import process of largely hydrophobic mitochondrial proteins, either targeted to the OMM directly (Jores et al., 2018) or the multiple membrane-spanning domain inserted into the IMM (Bhangoo et al., 2007).

A major argument for a post-translational mechanism of mitochondrial protein import has been the absence of a single specific ribosome-related regulatory and a targeting factor like the signal recognition particle in the case of protein import into the endoplasmic reticulum (Walter et al., 1981). However, co-translational mechanism cannot be excluded, which is heavily supported by the existing data. Early microscopic data exhibited the presence of ribosomes attached to the OMM (Kellems et al., 1975). Recently, polysomes directly attached to the TOM complex were observed *in vitro* by electron cryo-tomography (Gold et al., 2017). A co-translational import mechanism was shown for the enzyme fumarase with the isolated mitochondria (Knox et al., 1998). Fumarase was not imported into the organelle when expressed separately. In contrast, a significant amount of protein was detected inside the mitochondria when the fumarase preprotein was translated in the presence of mitochondria when the polypeptide is able to directly engage mitochondrial translocation machineries before the translation is completed or the protein is fully folded.

Both localized mRNA sequencing (Fazal et al., 2019) and localized ribosome profiling (Williams et al., 2014) show a large amount of tightly associated and actively translated mRNA near the OMM. Some mRNAs are attached via nascent-chain of actively translating ribosomes, and dissociates from the OMM in the absence of either ribosome translation or mitochondrial potential. Therefore, active ribosomes are associated with the translocation machinery by the interaction of the nascent polypeptide chain with the TOM complex. Additional indirect evidence for co-translational import comes from genetic studies. The deletion of OM14 (a tail-anchored OMM protein) leads to the reduced association of ribosomes with OMM (Lesnik et al., 2014) as well as Tom20 deletion causes partial mRNA dissociation from OMM (Eliyahu et al., 2010). Taken all these findings together, both co- and post-translational processes have certain roles in mitochondrial protein transport; however, it remains unclear which one is more important for the living cell. No experimental approach has been able to distinguish between those two mechanisms for living cells. Here we show that microinjection of exogenously produced proteins into living cells is a valuable tool in this respect.

Direct studies of mitochondrial protein import remain, however, largely limited. The usual approach to introduce protein into the cell is in the form of DNA via transfection followed by translation by cell machinery. It is not possible to study protein transport dynamics with this approach.

To reliably study the rate of protein transport inside living cells, the following requirements should be met: i) quick (compared to protein kinetics) delivery of protein into the cell, ii) low fluorescence background signal, iii) usability in the adherent cells allowing microscopic observations. The delivery of a recombinant protein into the cell can be achieved by various approaches, such as physical membrane disruption, protein modifications, or usage of nanocarriers (Du et al., 2018). Protein alterations (i.e., hypercharge, cell penetrating peptides, or poly-cysteine motifs) likely affect the protein behavior inside the cell, while carriers require complicated engineering to avoid endosomal entrapment. Additionally, the usage of protein in the vicinity of the cell, as is also the case with physical methods, such as electroporation and optoporation, generates a fluorescence background signal from the protein of interest and lacks a clear starting point.

Here we developed a procedure for a gentle microinjection of plasmids and recombinant proteins into living cells with further monitoring of protein import into the mitochondria by fluorescence microscopy inside living cells for up to 3 h. We show that our injection protocol is sufficiently non-invasive to preserve the mitochondria protein transport machinery and cell well-being. By injecting the fluorescent MTS-SNAP-tag protein, we show its mitochondrial import, which does not occur with the deletion of MTS or mitochondrial potential. When injecting into cells pretreated with translation inhibitors – puromycin (PUR) and cycloheximide (CHX) – we detected their different effects on the mitochondrial import rate. Our results imply that around 50 % of all TOM complexes are constantly occupied by co-translationally transported preproteins in a living cell at rest. We believe that microinjection is a valuable tool not only for the studies of mitochondrial protein import in knock-out systems, disease, or aging models, but also as a method to investigate protein transport dynamics in other cellular organelles.

## Results

### Microinjection of recombinant proteins into living cells

To study protein import into the mitochondria of living cells, we developed an experimental procedure for microinjection of a recombinant fluorescent proteins (FP) into living mammalian cells with a subsequent real-time microscopic detection of its cellular redistribution (see Fig. 1). To establish the non-invasiveness of the injection for the mitochondria and the cell functioning, firstly, we injected the expression vector pMC MTS-EmGFP (emerald green fluorescent protein), encoding EmGFP targeted to the mitochondrial matrix, into adherent HeLa cells cultivated in a glass-bottom dish. (Fig. 2A, Video S1). We used the MTS of cytochrome c oxidase subunit 8A (COX8A) to target FPs to the mitochondria. In such a way, we can monitor the cellular localization of FPs from the first minute up to several hours in the course of their intracellular expression. The mitochondria were labeled by Mitotracker Orange (MTOrange), which is spectrally well separated from MTS-EmGFP. After one hour of incubation, a sufficient amount of MTS-EmGFP was produced, and its fluorescence signal built up in the mitochondria with a perfect match to MTOrange localization. MTS-EmGFP fluorescence appeared at later time points in the cytosol as well, however, in a lesser amount. This effect is probably caused by an overexpression of MTS-EmGFP to very high levels, which the mitochondrial protein import machinery cannot handle properly. Additionally, we performed similar experiments with the pMC MTS-Dendra2 expression vector. For several cells in this case we observed cell divisions hours after the plasmid injection that was followed by a boost of MTS-Dendra2 expression localized in the mitochondria (Video S2). From these results we concluded that our injection procedure is sufficiently non-invasive: it does not disturb the mitochondrial health (as evidenced by the intact mitochondria shape and the functional protein import machinery) and the expression systems of the cell (as evidenced by the expression from the plasmid) in the course of the experiment. Even vital cell functions were not compromised as cell division observations for the pMC MTS-Dendra2 injection show.

**Figure 1.**
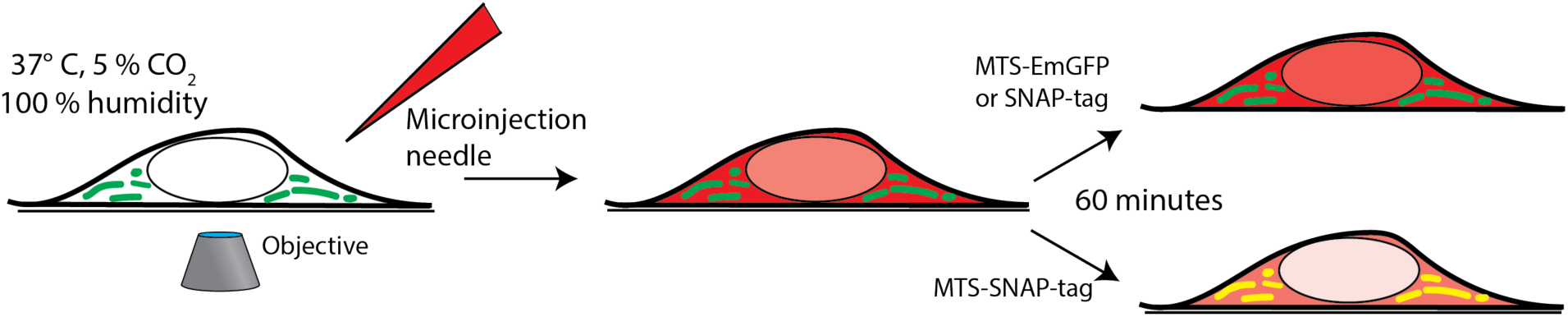
Experimental layout. Microinjection is performed on the cells in the field of view of an inverted microscope, and time series imaging is performed. The cells are grown in a 35 mm glass-bottom imaging dish and placed in an incubator mounted on an inverted confocal microscope. The microinjection of the protein (red) is performed, and co-localization (yellow) with the mitochondria (green) is tracked over time.

**Figure 2.**
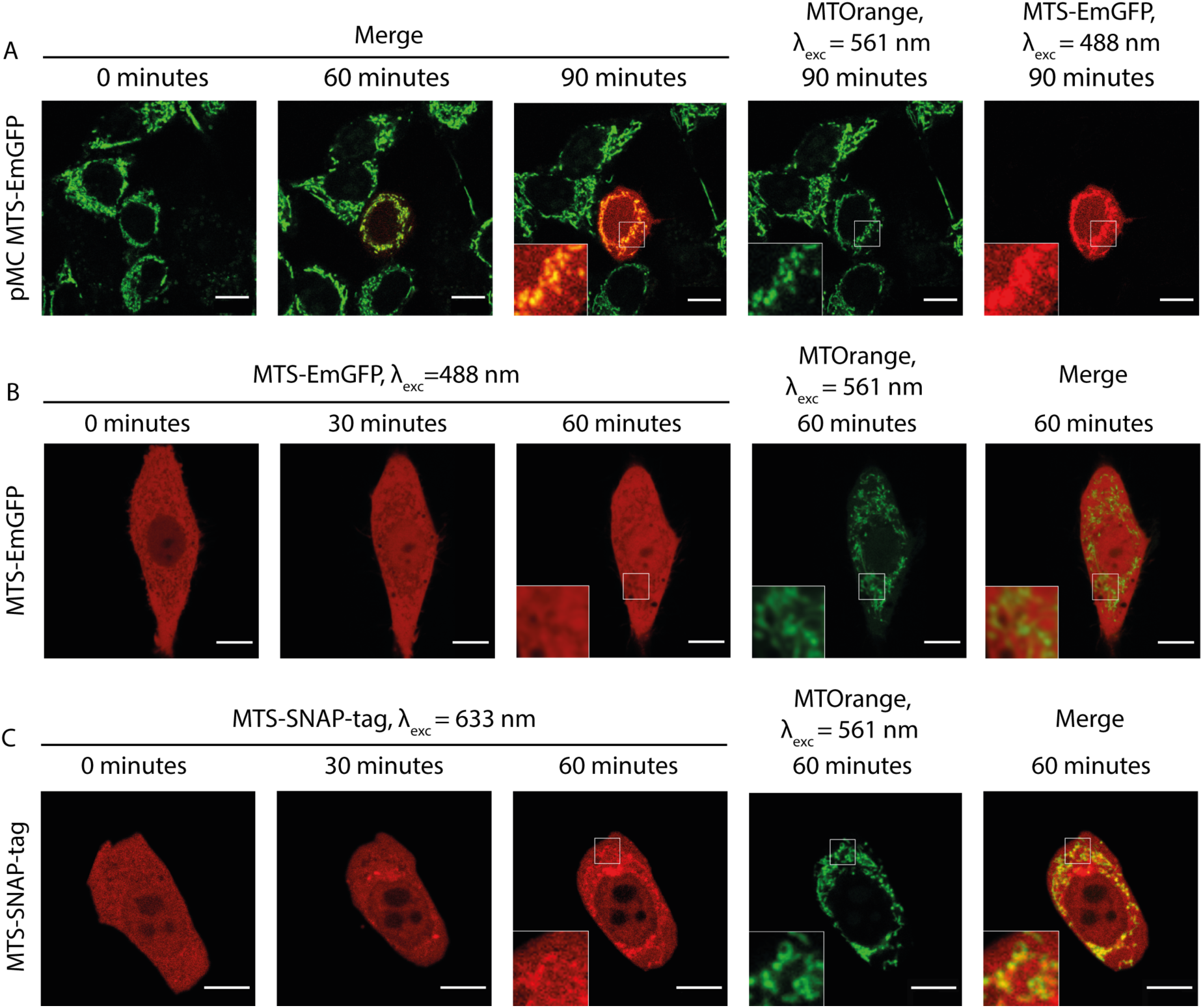
Microinjection in HeLa cells. (A) Time-lapse microscopy of the MTS-EmGFP (red) expression after the injection of the expression vector pMC-MTS-EmGFP in the HeLa cells. The MTS-EmGFP fluorescence is observed first at ca. 60 min and intensifies for several hours. The mitochondrial network labeled by MTOrange (green) appears normal at all times after the injection, and the newly synthesized and matured MTS-EmGFP localizes in the mitochondria. (B) Time-lapse microscopy of the injected MTS-EmGFP protein distribution in the HeLa cells. The MTS-EmGFP (red) evenly distributes inside the cytoplasm and nucleus, while the mitochondria labeled by MTOrange (green) are visible as regions with a lower fluorescence intensity (‘shadows’), which correlates well with the strong fluorescence at the corresponding pixels in the MTOrange image. (C) Time-lapse microscopy of the injected MTS-SNAP-tag protein distribution in the HeLa cells. The MTS-SNAP-tag (red) fluorescence shows structures with a higher fluorescence intensity correlating with the MTOrange labeled mitochondrial network (green) from 30 min after the injection, seen more clearly 60 min after the injection. Scale bars: 10 µm.

With the established experimental parameters for microinjection, we moved onto the injection of recombinant FP. FPs were expressed in *E. coli* as part of a chimeric protein (Fig. S1A). The FP core is N-terminally fused with MTS followed by the SUMO protein. The role of SUMO is to protect MTS from proteolytic degradation during expression and purification (Malakhov et al., 2004). The degradation is likely to be caused by the N-terminal pathway in *E. Coli* and the unspecific exoproteases activity inside bacterial cells (Gonzales and Robert-Baudouy, 1996). SUMO is cleaved by ULP1 SUMO protease after purification leaving intact MTS at the N-terminus of FP (Mossessova and Lima, 2000). As FPs, we used an EmGFP (Cubitt et al., 1999) and the SNAP-tag protein, a self-labeling variant of the human enzyme *O*^6^-alkylguanine DNA alkyltransferase (Juillerat et al., 2003).

We injected the purified recombinant MTS-EmGFP into adherent HeLa cells. As shown in Fig. 2B and Video S3, the injected MTS-EmGFP dispenses fast in the cytosol while equilibration with the nucleus is completed only after 30-60 min. The MTS-EmGFP does not enter the mitochondria for up to 2 h of the observation time, as can be clearly seen by low-intensity areas (later referred to as ‘shadows’). These ‘shadows’ overlap to a large extent with the MTOrange staining and, therefore, were identified as the mitochondria. The low but non-zero EmGFP-fluorescence observed in these mitochondrial regions was possibly caused by out-of-plane fluorescence from proteins in the cytoplasm surrounding the mitochondria. Some ‘shadows’ have no counterpart in the MTOrange staining, which most likely correspond to the endoplasmatic reticulum parts. The volume of the injected MTS-EmGFP solution can be estimated as 1-4 % of the cell volume (see the “Methods” section for details), which is in the range of naturally occurring cell volume regulation and does not harm the cell significantly. The mitochondrial network remained intact and morphologically unchanged in the course of the experiment.

The absence of the mitochondrial protein import unambiguously shows that no post-translational transport is possible for MTS-EmGFP. Presumably, the bulky and highly stable β-barrel structure of EmGFP precludes the post-translational import. To further explore this phenomenon, we performed import experiments using radiolabeled MTS-Dendra2, into the mitochondria isolated from yeast with a well-characterized experimental setup (Becker et al., 2009). Although another, highly efficient, MTS sequence was used (see Fig. S2 and the description therein), the same lack of import of MTS-Dendra2 with its very similar to the EmGFP three-dimensional structure was observed in the standard *in vitro* import assay.

As β barrel-type FPs were not suitable for studying protein transport in our experimental setup, we focused on a different fluorescent reporter protein, namely the SNAP-tag protein. A SNAP-tag moiety in the expression vector COX8A-SNAP-tag was previously used for labeling the mitochondria after cell transfection (Stephan et al., 2019). In the earlier study, it was also demonstrated that SNAP-tagged fluorescent reporter proteins could be efficiently imported into the isolated yeast mitochondria in an *in vitro* setup (Martin et al., 2006). The major advantage of the SNAP-domain in fusion proteins is its ability to self-label with almost any (functionalized) fluorophore at a single defined cysteine residue in the active center of the SNAP enzyme.

Similar to MTS-EmGFP, the MTS-SNAP-tag protein was expressed in *E. coli* as N-terminal fusion with the MTS and SUMO-protein (Fig. S1A and B) removed before the microinjection. The SUMO-MTS-SNAP-tag protein was labeled by SNAP-Rho14 (λ_exc_=633 nm; λ_em_=645 nm). The labeled MTS-SNAP-tag protein was injected into HeLa cells (Fig. 2C). Soon after the injection (< 2 min), the MTS-SNAP-tag protein was distributed homogeneously in the cytoplasm and with a lower concentration in the nucleus. 20-30 min after the injection, the MTS-SNAP-tag protein was concentrated in cytoplasmic structures most of which are identified as the mitochondria by the MTOrange staining (Fig. 2C, Video S4). The visible accumulation continues to rise even 2 h after the injection. However, we used a cutoff point of 90 min for the following experiments. In this time range, photobleaching, dynamic range limitations, and cellular movement do not interfere significantly with the observations.

In contrast to the MTS-EmGFP, the MTS-SNAP-tag protein is unambiguously imported into the mitochondria in a post-translational fashion. To the best of our knowledge, this is the first visualization of post-translational mitochondrial protein import observed in real-time in a living cell.

Although the Rho14 chromophore is considerably smaller compared to the SNAP-tag protein, we cannot exclude its influence on the protein import. Accordingly, we performed the experiments with other, commercially available, SNAP dyes: SNAP-Surface 594 and SNAP-Cell TMR-Star. The MTS-SNAP-tag protein labeled with either dye exhibited similar import into the mitochondria. However, a noticeable variation in the import time was observed, likely reflecting different labeling efficiency or possibly influence of the dye nature (Fig.S3 A, B). In all further experiments, we used Rho14 since it is spectrally well separated from green/yellow fluorescence of the mitochondrial markers used in this study: MTOrange and later MTS-Dendra2 (see below).

Our further improvement of the experimental setup is related to the mitochondria marker. The labeling efficiency of the available mitotracker dyes for living cells varies under experimental conditions and utilizes the IMM potential. For this reason, we created a HEK 293 cell line stably expressing MTS-Dendra2 with mitochondrial localization (see the “Methods” section for details of cell line generation). This approach provides a more convenient, stable, and cell-to-cell reproducible labeling of the mitochondria. In particular, this staining retains after the IMM potential depletion by Carbonyl Cyanide 3-ChloroPhenylhydrazone (CCCP), a widely used uncoupler of mitochondrial oxidative phosphorylation due to the inability of the MTS-Dendra2 to exit mitochondria in contrast to MTOrange (Fig. S4).

Microinjection of the fluorescent MTS-SNAP-tag protein into HEK 293-MTS-Dendra2 cells completely reproduce all import features observed in HeLa cells, demonstrating the suitability of a different cell type for mitochondrial protein import experiments in living cells (Fig. 3A, Video S5).

**Figure 3.**
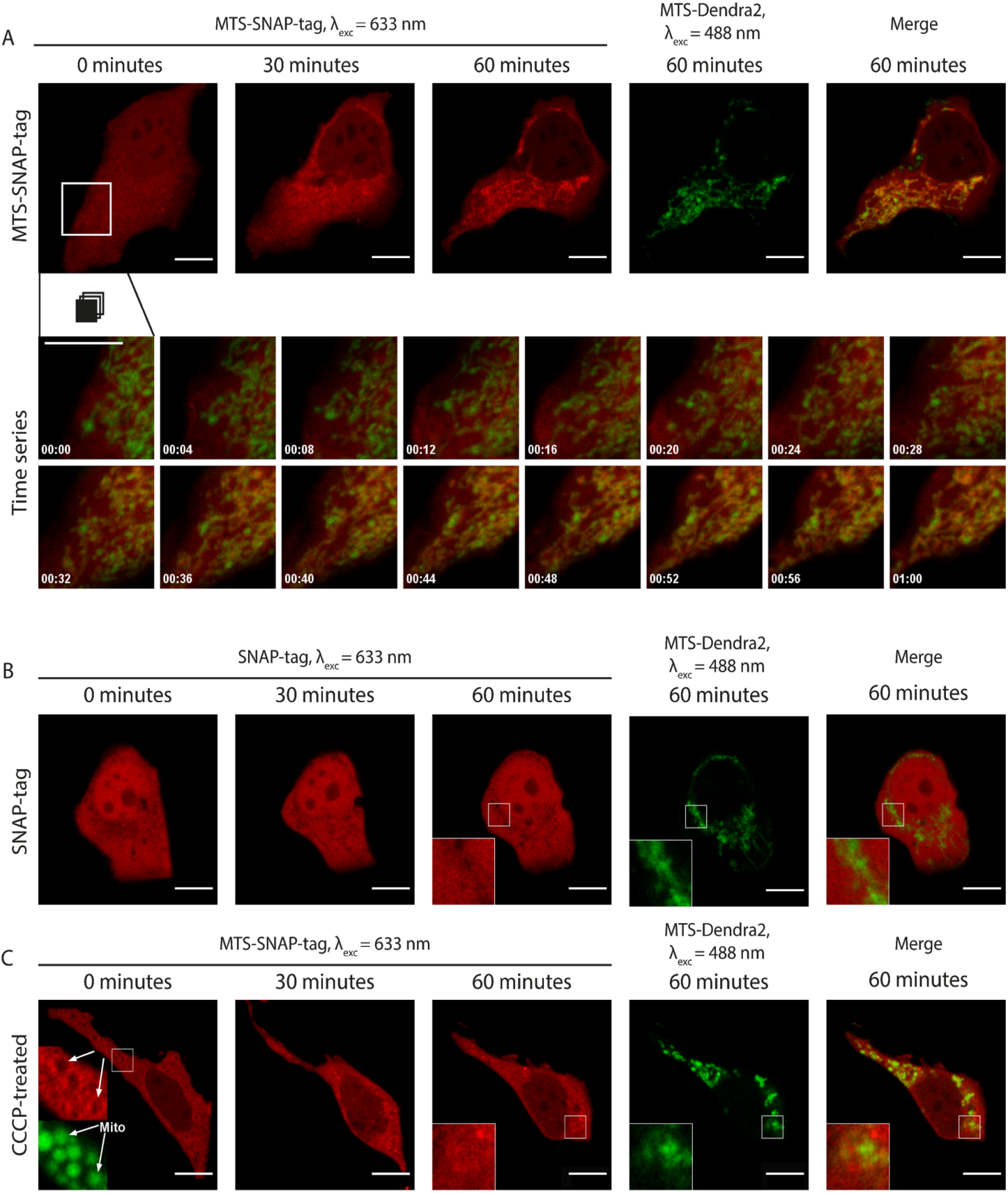
MTS-SNAP-tag protein import into the mitochondria. (A) Time-lapse microscopy of the MTS-SNAP-tag protein redistribution in the HEK 293-MTS-Dendra2 cells after the injection. The MTS-SNAP-tag protein fluorescence (red) co-localizes with the mitochondria after 30 min. The mitochondria are labeled by the MTS-Dendra2 protein (green). The bottom row shows a time series of the selected region. The tracking time is given in minutes. (B) Time-lapse microscopy of the SNAP-tag protein redistribution (red) in the HEK 293-MTS-Dendra2 cells after the injection, the ‘shadows’ correspond to the mitochondria, as seen in the enlarged inset by comparison with the MTS-Dendra2 labeled mitochondria (green). (C) Time-lapse microscopy of the MTS-SNAP-tag protein redistribution after the injection in the HEK 293-MTS-Dendra2 pre-incubated with 50 µM CCCP. The MTS-SNAP-tag protein (red) initially forms a brighter rim around the spherical mitochondria (green), shown enlarged in the insets in the first image (0 min). Scale bars: 10 µm.

The experimental setup described above using of HEK 293 with the MTS-Dendra2-labeled mitochondria and the injection of SNAP-tag labeled with Rho14 was used in all later experiments. To study the speed of mitochondrial protein import in living cells in each experiment, we collected confocal fluorescence images every 2 min after the injection typically for about 90-120 min after the injection. A time series of fluorescent images composes a ‘movie’ of the MTS-SNAP-tag protein transport into the mitochondria (see Fig. 3A, video S5). With this setup, one can clearly detect the import of the MTS-SNAP-tag protein into the mitochondria in the merged image by the disappearance of green and the concomitant appearance of yellow mitochondria at later time points, most pronounced between 20 and 50 min after the protein injection. The experiments described below were repeated, and all the described observations were reproduced for at least five cells.

To validate our experimental setup and to exclude potential artifacts of the protein purification and labeling, we produced the SNAP-tag protein without an MTS. The SNAP-tag protein was labeled with SNAP-Rho14 and injected into the cell (Fig. 3B, Video S6). Similarly to the MTS protein variant injection, the SNAP-tag protein was evenly distributed in the cytoplasm right after the injection. However, no change in its spatial distribution was observed during the next 90 min. Instead, similar to MTS-EmGFP, the mitochondria were seen as ‘shadows’ in the uniform SNAP-tag fluorescence in the cytoplasm. The ‘shadows’ coincide well with the MTS-Dendra2 fluorescence of the mitochondria (see insets in Fig. 3B). These ‘shadows’ are a direct consequence of the exclusion of the SNAP-tag protein from the mitochondria.

### Effects of the uncoupling of the IMM potential on the MTS-SNAP-tag protein import in living cells

The IMM potential is a major driving force for the import of mitochondrial preproteins. To assess the effects of uncoupling chemicals, dissipating the IMM potential, we injected the MTS-SNAP-tag protein into the cells pretreated with CCCP. Disrupting the mitochondrial potential, CCCP interferes with its tubular morphology (Legros et al., 2002). The mitochondria become spherical or elliptical, which we also observed in our experiment (Fig. 3C, Video S7). Right after the injection, the mitochondria appeared as ‘shadows’ in the MTS-SNAP-tag image with a bright rim around them (see inset in Fig. 3C at 0 min). It probably represents the MTS-SNAP-tag protein molecules that were attached to the mitochondrial surface via the TOM machinery but were unable to translocate through the IMM due to the absence of the potential. This feature disappears over time (in approx. 10 min). The MTS-SNAP-tag was not efficiently imported into the mitochondria as compared to untreated cells. However, the MTS-SNAP-tag protein ‘shadows’ were also visible to a much lesser extent in the CCCP-treated cells than in the experiment with the SNAP-tag w/o MTS injected in non-treated cells. Therefore, we cannot exclude the possibility that a low amount of protein import still can occur in the absence of the membrane potential. An alternative explanation is that due to the change in the mitochondrial morphology (spheres/ellipses instead of the mitochondrial network), the mitochondrial ‘shadows’ are less resolved and more smeared in the case of the CCCP-treated cells compared to non-treated ones. In summary, however, protein accumulation inside the mitochondria under these conditions is almost negligible compared to protein import levels in the untreated cells. Consequently, we conclude that the mitochondrial potential is necessary to efficiently import preproteins targeted to the IMM or matrix in living cells.

### Influence of the mRNA-ribosome-nascent chain complex (RNC) state on mitochondrial protein import in living cells

In our experimental setup, we use living cells that actively translate endogenous proteins, some of which are targeted to mitochondria. The injected reporter protein competes for mitochondrial import with the endogenous ones. We use PUR and CHX to prevent new protein production and consequently stop new protein import into the mitochondria. Both compounds are widely used translation inhibitors, but they work via different mechanisms. PUR causes premature translation termination and nascent chain release from the ribosome (Nathans, 1964), whereas CHX stalls translation but preserves the RNC (Ennis and Lubin, 1964). We injected the Rho14-labeled MTS-SNAP-tag protein into the HEK 293-MTS-Dendra2 cells pretreated for 30 min with either PUR or CHX. As shown in Fig. 4, in both cases, we clearly observed protein import much alike as in the untreated cells. However, the rates of import were significantly different between the two. The mitochondrial protein import was faster in the PUR-treated cells – the Rho14 fluorescent signal localized to the mitochondria was already clearly visible as early as 10 min after the injection (Fig. 4C, Video S9). CHX had almost no effect on the mitochondrial protein import rate compared to the untreated cells – the MTS-SNAP-tag protein import occurred on the time scale of 30-60 min (compare Fig. 4 B with A, and Video S8 with S5).

**Figure 4.**
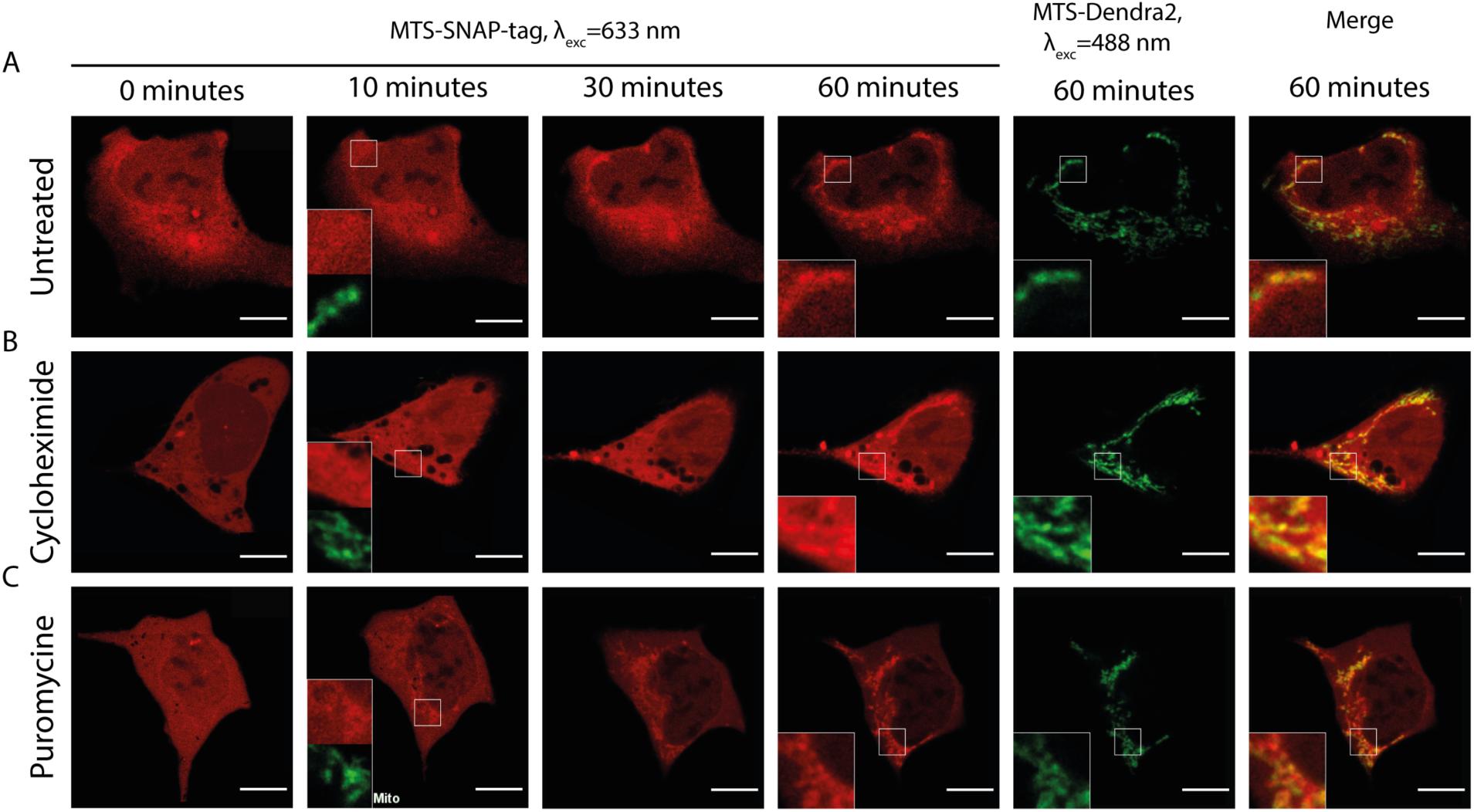
MTS-SNAP-tag protein import into the mitochondria under CHX and PUR treatments. Time-lapse microscopy of the microinjected MTS-SNAP-tag protein (red) import into the mitochondria in the HEK 293 MTS-Dendra2 (green) cells under different conditions. (A) The untreated HEK 293-MTS-Dendra2 cells. After 10 min, no noticeable MTS-SNAP-tag protein accumulation in the mitochondria can be seen (shown enlarged in the inset). MTS-SNAP-tag protein accumulation is seen after 60 min. (enlarged region in the insets). (B) The HEK 293-MTS-Dendra2 cell treated with 100 μg/ml CHX. After 10 min, no noticeable MTS-SNAP-tag protein accumulation in the mitochondria can be seen (shown enlarged in the inset). The MTS-SNAP-tag protein accumulation is seen similarly to the untreated cells (enlarged region in the insets) after 60 min. (C) The HEK 293-MTS-Dendra2 cell treated with 20 μg/ml PUR. After 10 min, the MTS-SNAP-tag protein accumulation in the mitochondria can be seen (shown enlarged in the inset). After 60 min, the MTS-SNAP-tag protein accumulation results in brighter mitochondria as compared to the CHX-treated and untreated cells. Scale bar 10 μm.

Next, we quantified the rate of the mitochondrial protein import for the untreated, PUR- and CHX-treated cells. For that, 7-9 cells were chosen for each condition from at least three different cell dishes with separate protein purification batches for each dish. We related the mitochondria contrast against cytosol (see the “Methods” section for details) to the amount of the imported protein. The increase in contrast with time is, therefore, proportional to the increase in the amount of the imported protein. Figure 5A depicts the contrast time-series averaged over all measured cells for a given condition. We treated our previously described results on the injection of SNAP-tag w/o MTS in the same way and added these data for comparison. We linearly fitted the measured mitochondrial contrast for 5-60 min and used the slope coefficient as a proxy measurement of the import rate. The import rate for each cell is plotted in a box chart in Fig. 5B. Consistent with the previous results, the import rate for the protein w/o MTS is close to zero. No differences in the mitochondrial protein import rates can be detected between the untreated and CHX-treated cells (p≈0.4); however, the import rate in the PUR-treated cells is ∼2-times faster compared to the CHX-treated or untreated cells (p≈0.01 and 0.03, respectively). As described in the discussion, the different import rate for the PUR- and CHX-treated cells is likely connected to the RNC attached to the TOM complexes on the cytoplasm side of the OMM.

**Figure 5.**
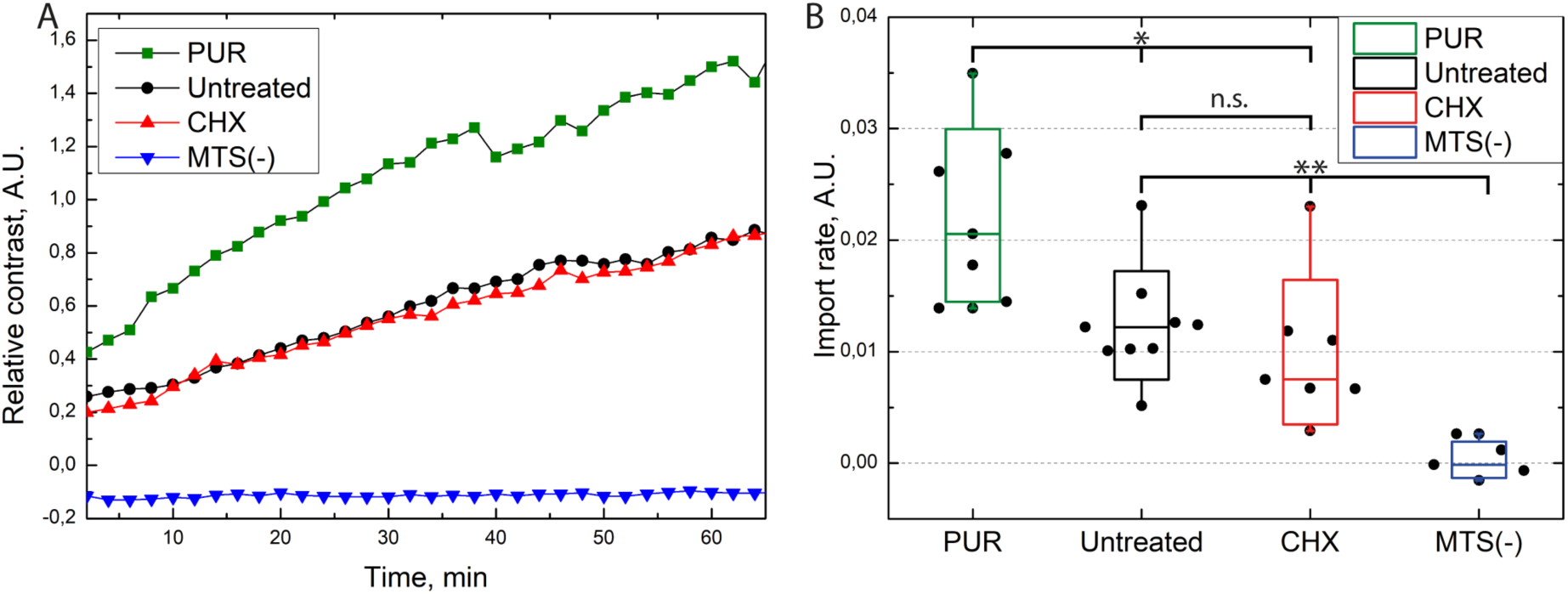
Import rate of the MTS-SNAP-tag protein into the mitochondria. (A) MTS-SNAP-tag protein import into the mitochondria. Each line represents mitochondrial contrast over time averaged for 6-9 cells under similar conditions. The PUR treated cells (green) show a higher import speed compared to the CHX- and untreated cells (red and black, respectively). The SNAP-tag protein (blue) shows no import into the mitochondria. (B) Import rate under different conditions. The contrast of the MTS-SNAP-tag protein imported into the mitochondria over time is linearly fitted between 5 to 60 min, and the slope coefficients are taken. The boxes show a standard deviation, the whiskers indicate the range of values. 7/7/9 cells from three separate experiments were used for puromycin/cycloheximide/no treatment, respectively. For the SNAP-tag protein experiment, 6 cells were used from a single experiment. The significance level is given according to the Mann-Whitney test: ** -p<0.005, * -p<0.05, n.s. -not significant.

### Effect of CHX and PUR on the mitochondrial protein import in the isolated mitochondria

To put the observations of the microinjection experiments into the context of the previous experimental work on the mitochondrial protein import, we also performed *in vitro* mitochondrial protein import experiments, which are typically based on a post-translational approach. Here, mitochondrial preproteins are synthesized in a cell-free system, in a radiolabeled form, and incubated with the isolated, intact, and energized mitochondria in an appropriate buffer system. We used the artificial reporter construct Su9-DHFR, consisting of a mitochondrial targeting sequence derived from the subunit 9 of the mitochondrial Fo-ATP synthase from the mold *Neurospora crassa* and the cytosolic dihydrofolate reductase (DHFR) enzyme from the mouse as a cargo moiety. This preprotein was synthesized and labeled with [S]^35^-Methionin by *in vitro* transcription and translation in rabbit reticulocyte lysate. The radiolabeled Su9-DHFR was efficiently imported *in vitro* into the mitochondria isolated from human HeLa cells, as expected (Fig. 6A). The analysis of the radioactive proteins associated with the mitochondria by SDS-PAGE and autoradiography indicated an increasing amount of the processed form (p) over time. The p-form is generated by the matrix processing peptidase, indicative of the removal of the targeting sequence after translocation into the mitochondrial matrix. In addition, the imported radiolabeled preproteins were protected against degradation by the externally added protease K (PK) due to the still intact mitochondrial membranes. Typically of a successful import reaction, both processing and acquisition of protease resistance are dependent on the presence of an intact IMM potential, as shown in the control experiments where the mitochondria were pretreated before import with inhibitors of oxidative phosphorylation and uncoupling chemicals (-Δψ). The post-translational import of Su9-DHFR under *in vitro* conditions appeared to be fast with detectable processing and translocation already in the time frame of minutes. A pretreatment of the isolated mitochondria by CHX or PUR, in order to affect potentially bound ribosomes that had not been removed by the isolation procedure, did not change the import efficiency or kinetics.

**Figure 6.**
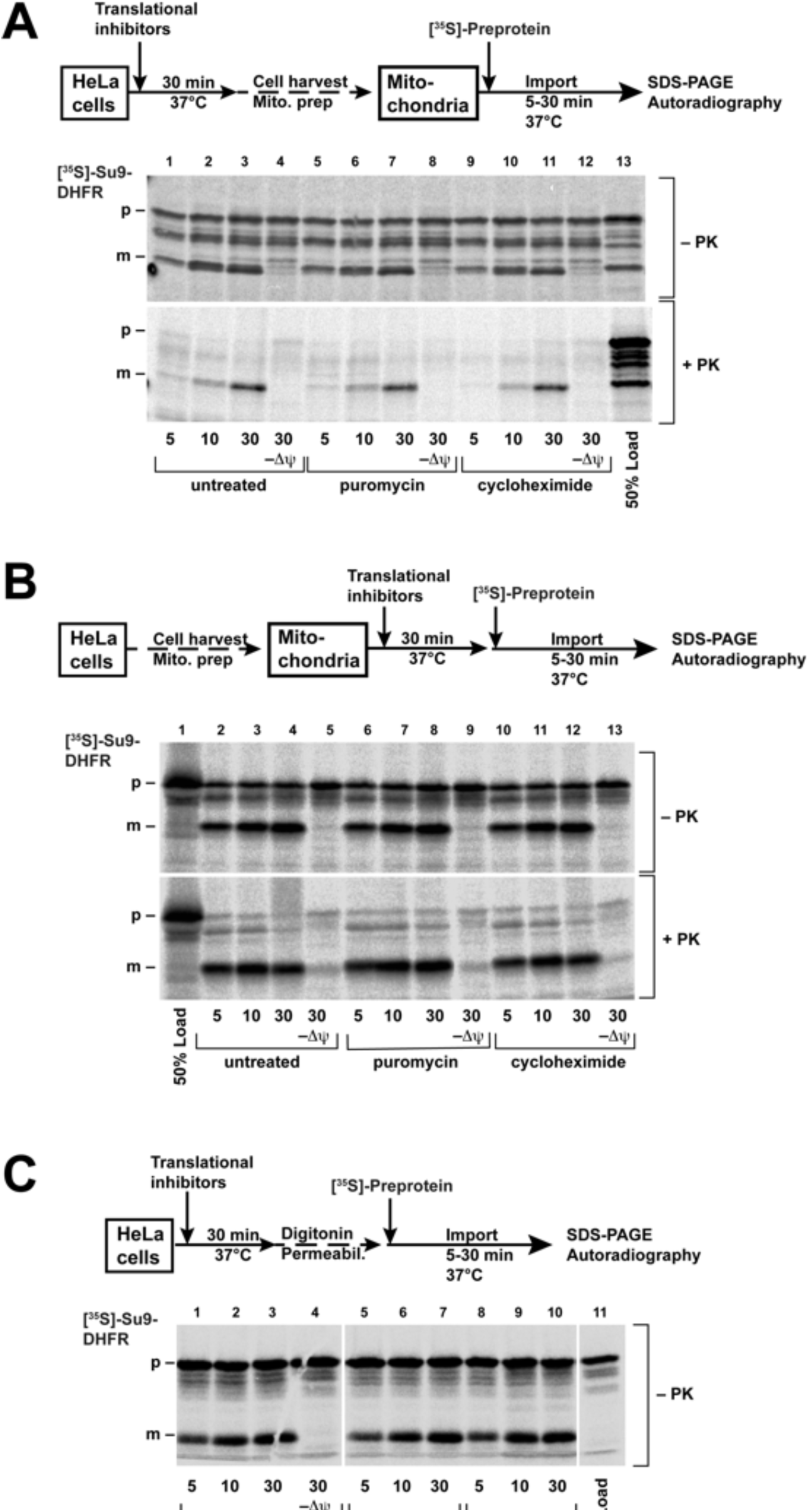
Import of the radiolabeled proteins into the mitochondria *in vitro*. Import of the [^35^S]-labeled protein Su9-DHFR was performed as described in the “Methods” section. The cells (A) or the isolated mitochondria (B) were pretreated before import with PUR (20 µg/ml) or CHX (100 µg/ml) for 30 min at 37°C. The import reactions were incubated for the indicated times. (C) The cells were permeabilized with digitonin after pretreatment, and then import was performed directly without isolation of the mitochondria. In the control reactions (-Δψ), the inner membrane potential was depleted, as described. After import, the cells were treated with proteinase K, as indicated. The imported proteins were analyzed by SDS-PAGE and autoradiography (p: precursor form, m: mature form of Su9-DHFR). The schematic outlines of the experimental procedures are given.

In order to most genuinely reproduce the conditions of the living cells, we also pretreated the intact HeLa cells in culture medium with the same amounts and times of CHX and PUR as were used in the injection experiments, isolated mitochondria from these cells, and repeated the import experiments using radiolabeled Su9-DHFR, as described above (Fig. 6B). Again, no difference between the import rates in the mitochondria isolated from control and the pretreated cells was observed.

We also developed a novel technical procedure where we used semi-permeabilized cells for mitochondrial protein import experiments. In this case, the elaborate mitochondria isolation procedure is circumvented, and any experimental manipulation of the cell system is reduced to a minimum. The permeabilization of the cell membrane was achieved by treatment of the cells harvested from the culture with the mild detergent digitonin under isotonic buffer conditions. The radiolabeled Su9-DHFR was added to this suspension of the permeabilized cells. After different incubation times, the cells were re-isolated, and the radioactive proteins were analyzed by SDS-PAGE and autoradiography (Fig. 6C). Similar to the traditional import experiments with purified mitochondria, we observed fast processing and also protease resistance of the Su9-DHFR preprotein, indicative of a successful and efficient mitochondrial protein import process. However, even under these conditions mimicking living cells, the presence of ribosomal inhibitors did not make a difference to the import reaction.

Taken together, these experiments indicate that in the *in vitro* setup of a mitochondrial protein import reaction, the post-translational import exhibits kinetics different from injection in living cells, since both translational inhibitors are unable to change the import reaction. In contrast, our novel approach demonstrates differences and thus, much closer resembles the genuine situation of mitochondrial protein import in living cells, particularly concerning translocation kinetics and the relationship between post- and co-translational import processes.

## Discussion

Current knowledge of protein import into mitochondria originates mainly from *in vitro* experiments on the isolated intact mitochondria supplied with exogenously expressed precursor proteins. On a cellular level, information on the mitochondrial protein import reaction relied mainly on the observation of an accumulation of endogenously expressed mitochondrial proteins in the cytoplasm. As maturation is only possible in the mitochondrial matrix, these proteins remain in their precursor form. The accumulation of precursor proteins can typically be observed only after treatments with mitochondrial inhibitors or severely affected mutants. In particular, under pathological situations, the traditional setup severely limits the amount and the quality of information obtained on mitochondrial protein import efficiency. Direct visual observation of mitochondrial protein import in living cells was impossible for a long time due to the lack of adequate approaches. Importantly, all previous studies of post-translational mitochondrial protein import were done on the mitochondria isolated from cells or tissues. Under these conditions, they lose their natural environment (cytosolic cofactors, involvement of the RNCs, microtubules, and other organelles) and also their native morphology, being transformed from a tubular network to separate spherical, double-bilayer vesicles. Although most principal processes – like oxidative phosphorylation – are preserved in the isolated mitochondria, some of their functional properties, including protein import, are likely to be compromised.

Here, we developed a setup for a direct investigation of transport based on confocal fluorescence microscopy and microinjection of pre-expressed fluorescent proteins into living cells. In our study, we preserved the natural state of the cell, including functional expression machinery and undisturbed mitochondria in their natural state. To enable fluorescent detection of the import, we used different FPs N-terminally tagged with an MTS as a model for putative protein targeted to the mitochondria. The use of an MTS for targeted delivery of the desired proteins to the mitochondria is widely applied to deliver fluorescent proteins for mitochondria labeling or to influence mitochondrial function (Hoffmann et al., 1994; Stephan et al., 2019; Tkatch et al., 2017).

In our first approach, we used the EmGFP as a reporter. However, the purified MTS-EmGFP showed no mitochondrial protein import after microinjection. A plausible reason is that the folded β-barrel cannot fit through Tom40 pore in the OMM – the β-barrel diameter is ∼30 Å, whereas the internal diameter of the Tom40 pore is ∼15 Å. At the same time the unfolding of the EmGFP during transport is hampered by the known high stability of the FPs β-barrel fold (Stepanenko et al., 2013). It should also be noted that while many experiments were performed with protein import into the isolated mitochondria, no successful import into the inner mitochondrial membrane or the mitochondrial matrix was ever shown for proteins with a stable β-barrel structure. Therefore, we switched to the SNAP-tag protein – an engineered variant of the enzyme O6-alkylguanine-DNA alkyltransferase, which is able to spontaneously form a covalent bond to virtually any fluorescent dye chemically fused with a benzylguanine. The purified and labeled MTS-SNAP-tag reporter showed successful import into the mitochondria in living cells. The SNAP-tag size is close to the EmGFP (23 kDa and 27 kDa, respectively) and also will not fit through the Tom40 pore in its folded state. However, the SNAP-tag has α+β-fold and is apparently more amenable to unfolding and transport through the Tom40 pore.

The mitochondrial uncoupling agent CCCP is known to stop the import of proteins into the isolated mitochondria (Schleyer et al., 1982) and is widely used as an important negative control in mitochondrial protein import experiments. Consistent with that, the CCCP-treated cells showed no noticeable protein import into the mitochondria in our experiments. It should be noted that in the presence of CCCP, the MTS-containing proteins can still interact with the import machinery of the OMM but will not be inserted by the translocase of the inner mitochondrial membrane (TIM). Hence, the MTS-SNAP-tag protein is initially recognized by the TIM-TOM machinery, which, however, lacks the driving force of the mitochondrial potential. Consequently, the polypeptides cannot be transported through the membrane, resulting in a higher concentration on the surface without any localization inside, seen as an initial brighter rim.

PUR and CHX are two commonly used translation inhibitors. Translation inhibition influences all protein production, including the TIM-TOM complexes themselves. However, as the pre-treatment exposure for 20 min was very short compared to endogenous protein expression rates, the influence on the amounts of the TIM-TOM components or other mitochondrial proteins is negligible. Pre-treatment of the cells with these inhibitors before the injection of the protein caused remarkably different effects on the mitochondrial protein import reaction – CHX did not change the import rate significantly, whereas PUR increased it approximately two times. It sheds light on the long-standing question of whether post- or co-translational import prevails for mitochondria-targeted proteins in living cells. The translational inhibitors affect the internal expression of cellular proteins, including those of endogenous mitochondrial proteins. CHX ‘freezes’ RNC attached to the OMM (Gold et al., 2017; Kellems et al., 1974) and, thus, clogs the TIM-TOM complexes. On the other hand, PUR dissociates RNCs and also releases them from the mitochondrial surface (Fazal et al., 2019), if they interact via a newly translated protein. Hence, we conclude that the observed 2-fold increased import rate of the injected proteins results from the removal of RNC attached to the TOM complexes by PUR, which became available for import. Therefore, about 50% of all TIM-TOM complexes were occupied by co-translational import of the endogenous proteins. In living cells, post-translational and co-translational import of endogenous proteins compete with each other, and the rate of post-translational import is strongly affected by the occupation of the TIM-TOM complexes with RNC.

Our work provides an insight into mitochondrial protein import inside living cells, directly demonstrating not only import of exogenous proteins into mitochondria, but indirectly demonstrating the significant role of co-translational mitochondrial protein import. Our approach to study mitochondrial protein import provides a valuable tool to study protein import directly in living cell systems under different conditions, such as import machinery knock-out studies or in model systems of cardiovascular and neurodegenerative diseases prominently involving mitochondrial defects (Cenini et al., 2016a; Pickrell and Youle, 2015).

## Acknowledgements

We thank M. Ergasheva for help with cell line handling, I. Melnikov for preliminary microinjection experiments. This work was supported by RFBR (№20-54-12010). V.B., A.M., V.G. and I.O. are supported by the Ministry of Science and Higher Education of the Russian Federation (agreement # 075-00337-20-03, project FSMG -2020-0003). J.H. and J.A. acknowledges DFG (grant numbers HA2868/14-1, SFB1116 A04 and grant numbers AL288/5-1, SFB1116 A04, respectively).

## Author contributions

G.B., T.G., V.B., W.V., J.H. and J.A. designed experiments. A.B. produced proteins, performed microinjection experiments, fluorescence microscopy and data processing. I.O., N.M., A.K. and A.S. produced plasmids and helped with protein expression. F.v.A., I.M., P.J. J.A. and J.H. generated HEK293 MTS-Dendra2 cell line. I.M. and D.B. helped with cell line maintenance, fluorescence microscopy, and data processing. G.B., V.G., V.B., T.G., J.H., J.A. and A.M. acquired funding. T.G. and V.B. supervised the project. A.B., V.B., T.G. analyzed data and wrote manuscript with the impact from W.V., J.H., J.A., G.B., V.G., A.M., I.O., A.K., I.M. and N.M.

## Materials and methods

### Plasmid preparation

pMC MTS-EmGFP and pMC MTS-Dendra2 plasmid were produced from pMC plasmid containing an MTS by restriction and insertion of the EmGFP and Dendra2. The gene of *S. cerevisiae* SUMO was composed of *E. coli* class II codons (Hénaut and Danchin, 1996) with the use of the DNABuilder software (The University of Texas Southwestern Medical Center at Dallas, USA). The nucleotide sequence at the 5’-end was additionally optimized using the RNA WebServer (Institute for Theoretical Chemistry, University of Vienna) in order to reduce the probability of RNA hairpin formation. The designed gene was synthesized by PCR from overlapping oligonucleotides (Stemmer et al., 1995) designed by means of DNAWorks (Hoover and Lubkowski, 2002). The synthesis of oligonucleotides was purchased at Eurogen JCS (Moscow, Russia). The SUMO-tag was fused via overlap extension PCR with the MTS-EmGFP gene. The resulting SUMO-MTS-EmGFP PCR fragment was added via ligation into the Pet15b expression vector between the XbaI/XhoI sites. To obtain plasmid with the MTS-SNAP-tag, the EmGFP fragment was exchanged by the AgeI/XhoI restriction followed by ligation of the SNAP-tag PCR fragment, resulting in SUMO-MTS-SNAP-tag chimera. The SUMO-SNAP-tag was also produced with overlap extension PCR and the following ligation into Pet15b plasmid between the XbaI/XhoI sites. All plasmids were verified by DNA sequencing for correct gene presence.

### Protein expression

Transformed with the expression vector Pet15b containing either SUMO-MTS-EmGFP, SUMO-MTS-SNAP-tag or SUMO-SNAP-tag *E. coli* cells were grown in lysogeny broth (LB) medium up to OD_600_∼1.0 and induced with 1mM IPTG (Helicon, Moscow, Russia). The cells were harvested after 3 h, centrifuged at 5000 g for 10 min, and the pellet was frozen at -80° C. The cell lysis was performed either by a microfluidizer or ultrasound (depending on the amount of cells), in the lysis buffer – 300 mM NaCl 50 mM NaH_2_PO_4_ pH=7. The lysate was centrifuged at 10000 g for 1 h, and the supernatant was applied onto affinity resin Ni-NTA (Qiagen, Dusseldorf, Germany) in a column. After washing the column with the lysis buffer, the protein was eluted with 200 mM imidazole dissolved in the lysis buffer. Imidazole was removed by dialysis against the cell buffer: 130 mM KCl, 10 mM NaCl, 2mM CaCl_2_, 20 mM NaHCO_3_ pH=7.2. After the dialysis, the solution was filtered through a 0.22 μm syringe filter (Millipore, Darmstadt, Germany) and stored at 4° C.

### Label preparation

BG-NH_2_ (New England Biolabs Inc., Ipswich, MA, USA) and NHS-Rho14 (ATTO-TEC GmbH, Siegen, Germany) were dissolved in DMF and mixed in 1:1 molar ratio in DMF, and 5x excess of triethylamine was added according to the manufacturer’s instructions. The reaction was performed overnight at 30° C. The purification was performed on SiO_2_ G-60 (Merck, Darmstadt, Germany) column with 50% DMF washing step and an elution with pure DMF, with functionalized SNAP dye being less soluble and slowly eluting in DMF. The dye was concentrated by evaporation in Centrivac (LabConco, MO, USA) using a vacuum pump ChemStar Dry (Gardner Denver, Milwaukee, WI, USA). The Surface-SNAP-ATTO594 and the SNAP-TMR-STAR dyes were obtained from New England Biolabs Inc (USA).

### Protein labeling and purification

The dyes were added to purified 6His-SUMO-MTS-SNAP-tag protein in 1:1 molar ratio and left for 1 h at 30° C. Them, the protein solution was centrifuged to remove the precipitates and the non-bound dye was removed by a solution exchange through a 30 kDa filter. The last step was repeated 3-5 times. After all the non-bound dye was removed, the His-tagged protease ULP-1 was added at a ratio of 1:500 to the SUMO-MTS-SNAP-tag protein (concentration measured by OD_280 nm_) for 1 h at 30° C. Afterward, the protein was again centrifuged to remove the precipitation and passed through the Ni-NTA column, with the MTS-SNAP-tag protein flowing through freely and 6His-SUMO sorbing. The flow through was collected, concentrated in 10 kDa columns, the protein concentration and the labeling efficiency were determined by spectrophotometry using the ratio of OD_label_/OD_280 nm_. The labeled protein was stored at 4° C for up to 2 weeks. The same procedure was followed for the SNAP-tag protein (without an MTS) and the MTS-EmGFP. The labeling procedure was skipped for the last one. The protein quality was controlled by SDS PAGE (see Fig. S1B).

### Microinjection

Microneedles were prepared from capillaries with an outer/inner diameter of 1.2/0.94 mm (Harvard Apparatus, Cambridge, UK) in Sutter P-2000 (Sutter instruments, Novato, CA, USA). The protein before the injection was centrifuged at 20 000 g for 1 h to remove small aggregates. The protein solution was back loaded into the needle, and the needle was installed into the micromanipulator InjectMan (Eppendorf, Hamburg, Germany). The micromanipulator was installed on the microscope allowing for immediate cell microscopy after the injection and was connected to the microinjector FemtoJet (Eppendorf, Germany). The microinjection was performed at 20-30 hPa injection pressure, 0.1 s injection time, and a compensatory pressure of 20-25 hPa. The preliminary experiments were performed to estimate the injection volume. The fluorescence intensity of the diluted protein solution was measured using the same microscopy settings (laser power, gain, objective) as with the injection experiments. At around the 50x dilution intensity was comparable to the median intensity in the cells, thus the injected volume was estimated to be 1 to 4% of the cell volume fluctuating between the cells.

### Cell culture

HeLa and HeK 293 MTS-Dendra2 Cells were grown in 25 cm^2^ flasks (Corning, Flintshire, UK) in DMEM (Gibco, Waltham, MA, USA) with 10% FBS (Gibco, USA) and PenStrep antibiotic (Gibco, USA). For the microscopy experiments, the cells were grown in a 35 mm glass-bottom imaging dish (Ibidi, Graefeling, Germany) to 40-60 % density. If additional reagents were used, they were added 20 min beforehand to reach the following concentrations: PUR (Applichem, Darmstadt, Germany) 20 μg/ml, (Applichem, Germany) 100 μg/ml, CCCP (Sigma Aldrich, Darmstadt, Germany) – 50 μm. The MTOrange (MitoTracker™ Orange CM-H_2_TMRos, Thermo Fisher Scientific, Waltham, MA, USA) staining in the HeLa cells was performed with 100-200 nM concentration for 2-3 min. MTGreen (MitoTracker™ Green FM, Thermo Fisher Scientific, USA) staining was performed with 500 nM concentration for 15 min. For the HEK 293 MTS-Dendra2 cells no additional staining was used.

### HEK-Dendra2 line generation

#### Cloning of a lentiviral expression vector for Dendra2 targeted to the mitochondria

The plasmid pCMV/myc/mito cut with Sal I and Not I to retain the MTS from subunit VIII of human cytochrome c oxidase and the C-terminal myc epitope tag as vector backbone. The coding sequence for Dendra2 was amplified from a plasmid clone and inserted into this backbone using the Gibson Assembly® Cloning Kit (New England Biolabs, Frankfurt, Germany); the primer sequences are available upon request. The Dendra2 coding region in the resulting plasmid was verified by DNA sequencing to exclude point mutations due to nucleotide misincorporations during the amplification procedure. From this plasmid, part of the cytomegalovirus immediate early promoter and the complete MTS-Dendra2-myc coding region were excised with Nde I and Xba I and transferred to the pLenti-FLAG-Trx-1 (Goy et al., 2014) cut with the same restriction enzymes to generate pLenti-MTS-Dendra2-myc.

#### Generation of a HEK 293 cell clone stably expressing Dendra2 targeted to the mitochondria

The 5×10^4^ HEK 293 cells were seeded on a 35mm tissue culture dish and transduced with the MTS-Dendra2-myc lentiviral particles, as previously described (Goy et al., 2014), using a multiplicity of infection of approximately 10. Starting six days after the transduction, the cells were subjected to selection with 5 μg/ml puromycin until all the cells in a non-transduced control were dead. During the selection procedure, the cells were kept in a 1:1 mixture of the complete growth medium and the conditioned medium from an exponentially growing HEK 293 culture. Single clones of the transduced cells were obtained by a limited dilution procedure (McFarland, 2000). Therefore, the transduced, selected cells were seeded in a 96-well tissue culture plate, with an average of 0.5 cells per well, in the same medium mixture as before. The successfully growing clones were analyzed by flow cytometry to assess the monoclonal origin, and the exclusive mitochondrial localization of MTS-Dendra2 was verified by fluorescence microscopy. One of the clones was used for all further studies.

### Fluorescence microscopy

The fluorescence microscopy was performed on an inverted laser-scanning confocal fluorescence microscope based on LSM780 (Zeiss, Jena, Germany). The 35 mm glass-bottom imaging dishes were kept in an incubator maintaining 37° C, 5% CO_2_, 100% humidity (Tokai Hit, Shizuoka, Japan) that was mounted on the microscope stage. After the injection procedure, time-series imaging was performed in confocal fluorescent microscopy λ-mode using a 34-channel QUASAR detector (Zeiss, Germany) set to the appropriate spectral range depending on the dyes. For excitation, a 488 nm or 561 nm laser was used simultaneously with a 633 nm laser to excite the MTS-Dendra2 or MTOrange and the SNAP-Rho14. The injection experiments of the MTS-EmGFP with the MTOrange labeling were done with 488 nm and 561 nm laser excitation. All the experiments were conducted with 1024×1024 (141×141 μm) image size using 100x (NA=1.46, oil immersive) objective. The autofocus was performed to the same z-plane using laser reflection on the sample dish glass surface before every image was taken. Afterwards spectral unmixing was performed in the ZEN software (Zeiss, Germany) using saved spectra from the non-injected cells (MTOrange or MTS-Dendra2) and a pure protein before the injection (MTS-EmGFP or MTS-SNAP-tag protein), respectively.

### *In vitro* import reactions

Import into the isolated mitochondria was essentially performed, as described (Cenini et al., 2016b). Briefly, the intact mitochondria were isolated from the cultured HeLa cells by differential centrifugation under isotonic buffer conditions. The radiolabeled preprotein Su9-DHFR was generated by *in vitro* transcription and translation in reticulocyte lysate in the presence of ^35^[S]-methionine. The radiolabeled preproteins were added to the isolated energized mitochondria (25 µg total mitochondrial protein per lane) and incubated for up to 30 min at 30° C. To assess complete translocation, the import reactions were divided, and one half was treated with 50 µg/ml proteinase K (Merck, Darmstadt, Germany) to remove all non-imported preproteins. The mitochondria were then re-isolated, and their protein content was analyzed by SDS-PAGE and autoradiography. For import into semi-intact whole cells, the cultured HeLa cells were harvested (0.25 million cells per lane) and treated with 0,005 % digitonin for 5 min at 25° C in the import buffer (250 mM sucrose, 20 mM HEPES pH 7.6, 80 mM KOAc, 5 mM Mg(OAc)_2_. The permeabilized cells were re-isolated by centrifugation for 10 min at 12000 g at 4° C and gently resuspended in the import buffer containing 5 mM glutamate, 5 mM malate, 1 mM DTT, and 10 mM K_3_PO_4_. The import reaction was started immediately by the addition of the radiolabeled preprotein, as described above. The pretreatments with translational inhibitors puromycin (f. c. 20 µg/ml) and cycloheximide (f. c. 100 µg/ml) (Merck, Darmstadt, Germany) were performed after harvesting of the cells in the import buffer for 30 min at 37°C.

### Data processing

The fluorescent images were manually segmented in FIJI (Schindelin et al., 2012) by a polygon tool into the regions with single, separated cells. In individual cells, a mitochondrial mask was created from the image of the used mitochondrial marker (MTS-Dendra2 or MTOrange). Due to imperfect masking and out-of-plane fluorescence, some of the cytosolic fluorescence from cytosol contributes to the mitochondrial one. The difference between the mitochondrial and cytosol average fluorescence intensity represents the amount of the imported protein. The intensity difference was normalized to an average intensity from the cytoplasm to correct for photobleaching, resulting in a ratio resembling the local contrast of the mitochondria. This ratio is referred in the text as the local contrast of the mitochondria and is proportional to the amount of the imported Rho14-MTS-SNAP-tag protein into mitochondria. The time-series of the contrast shown in Fig. 5A were linearly fitted in a 5-60 min time range for each cell, and the slopes were plotted in a box chart, as shown in Fig. 5B.

## SUPPLEMENTARY

**Figure S1.**
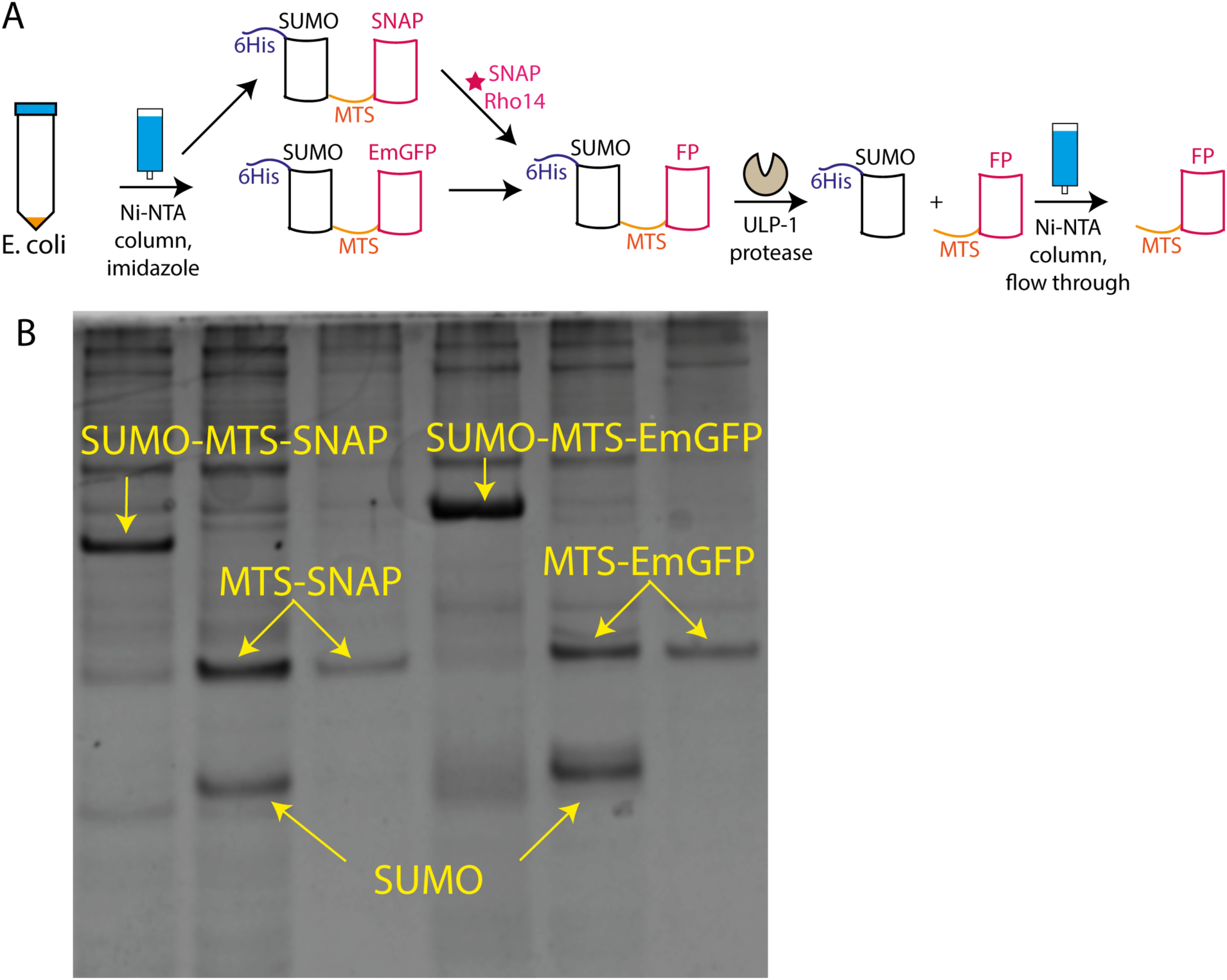
Protein expression. (A) Scheme of FP preparation. The initial purification by Ni-NTA is driven by 6His-tag contained in the SUMO protein. After purification (and labeling in the case of the SNAP-tag) SUMO is cleaved by His-tagged protease ULP-1, and the target protein flows through the Ni-NTA column, leaving the SUMO protein and protease on a column. (B) SDS page of the expressed proteins: the SUMO-MTS-SNAP-tag protein after initial purification (1^st^ column), mix of the SUMO and MTS-SNAP-tag proteins after ULP-1 cleavage (2^nd^ column), and the MTS-SNAP-tag protein after final purification (3^rd^ column). The three most right columns depict similar steps for the MTS-EmGFP purification.

**Figure S2.**
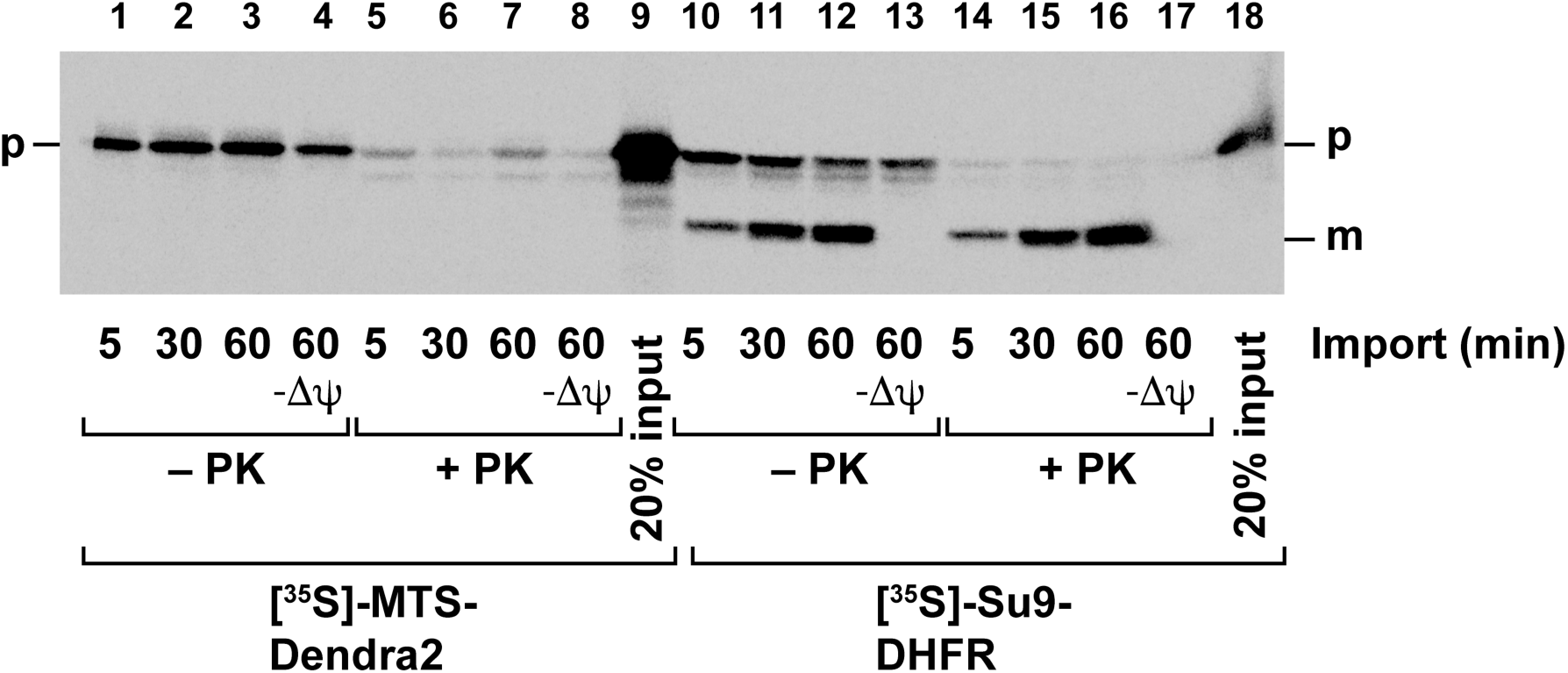
Import of the radiolabeled proteins MTS-Dendra2 and Su9-DHFR into the isolated yeast mitochondria. ^35^S-labeled proteins were generated by *in vitro* transcription/translation in reticulocyte lysate and stored at -80° C. The import reactions were performed essentially as described (Becker et al., 2009). The radiolabeled proteins were thawed and added to the intact and energized mitochondria isolated from the yeast cells. The import reactions were performed at 30° C for the indicated times. In control reactions (-Δψ), the mitochondrial inner membrane potential was depleted. The successful import reactions are characterized by processing of the added precursor (p) form to the mature (m) protein and its resistance against digestion by externally added proteinase K (PK). Both processing and protease resistance do not occur in the absence of a membrane potential. The Su9-DHFR was used as a positive control showing import to the mitochondria.

**Figure S3.**
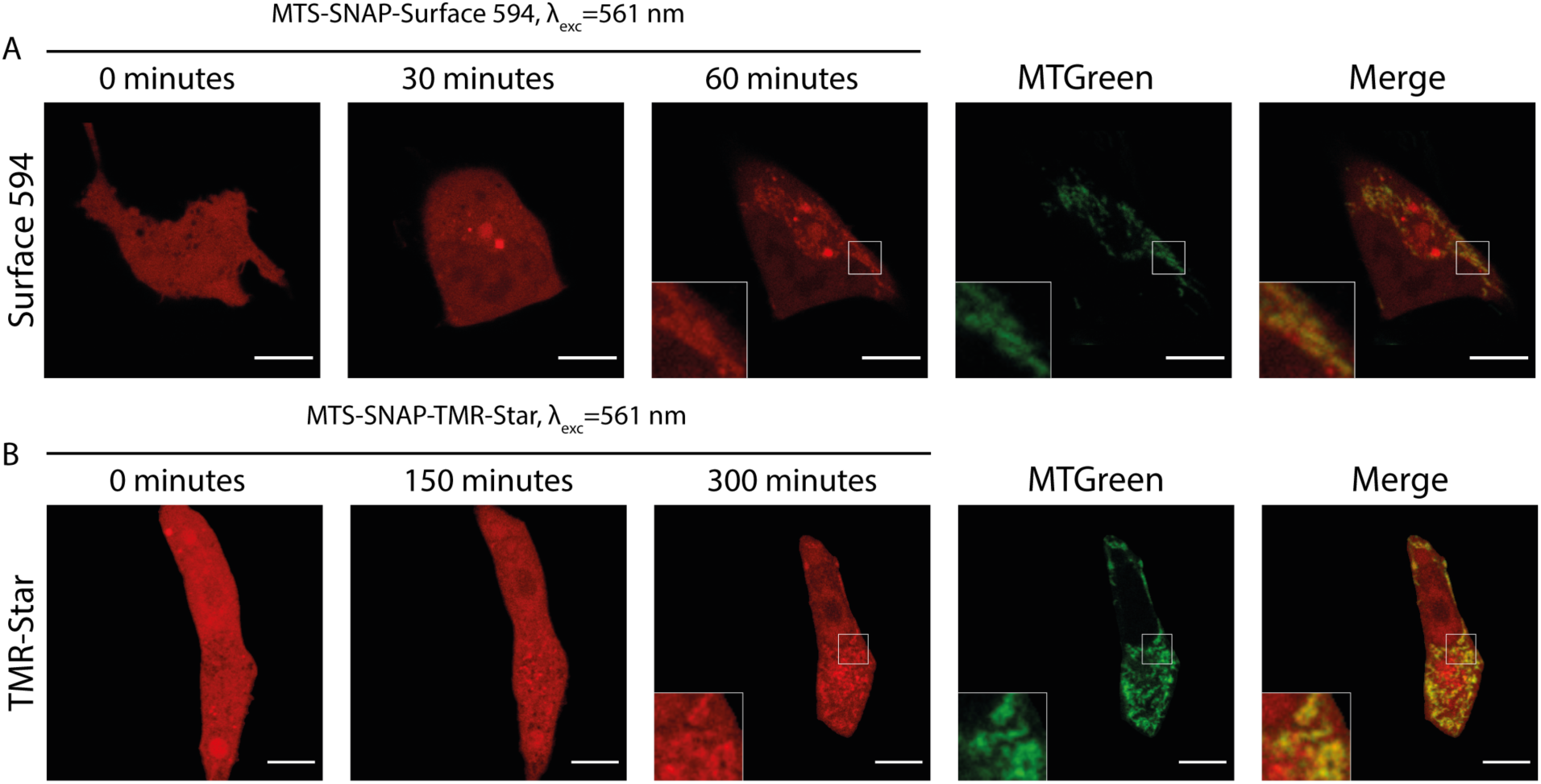
Import of the MTS-SNAP-tag protein labeled with commercially available dyes functionalized for binding to the SNAP-tag protein. Time-lapse microscopy of the microinjected MTS-SNAP-tag protein (red) import into the mitochondria labeled with the SNAP-Surface594 (A) and SNAP-TMR-Star (B). The HeLa cells mitochondria are stained with MTGreen (green). The SNAP-Surface-594 labeled MTS-SNAP-tag protein is imported with similar characteristic time to the Rho14 labeled protein, while the SNAP-TMR-Star labeled one is imported noticeably slower. Scale bar 10 μm.

**Figure S4.**
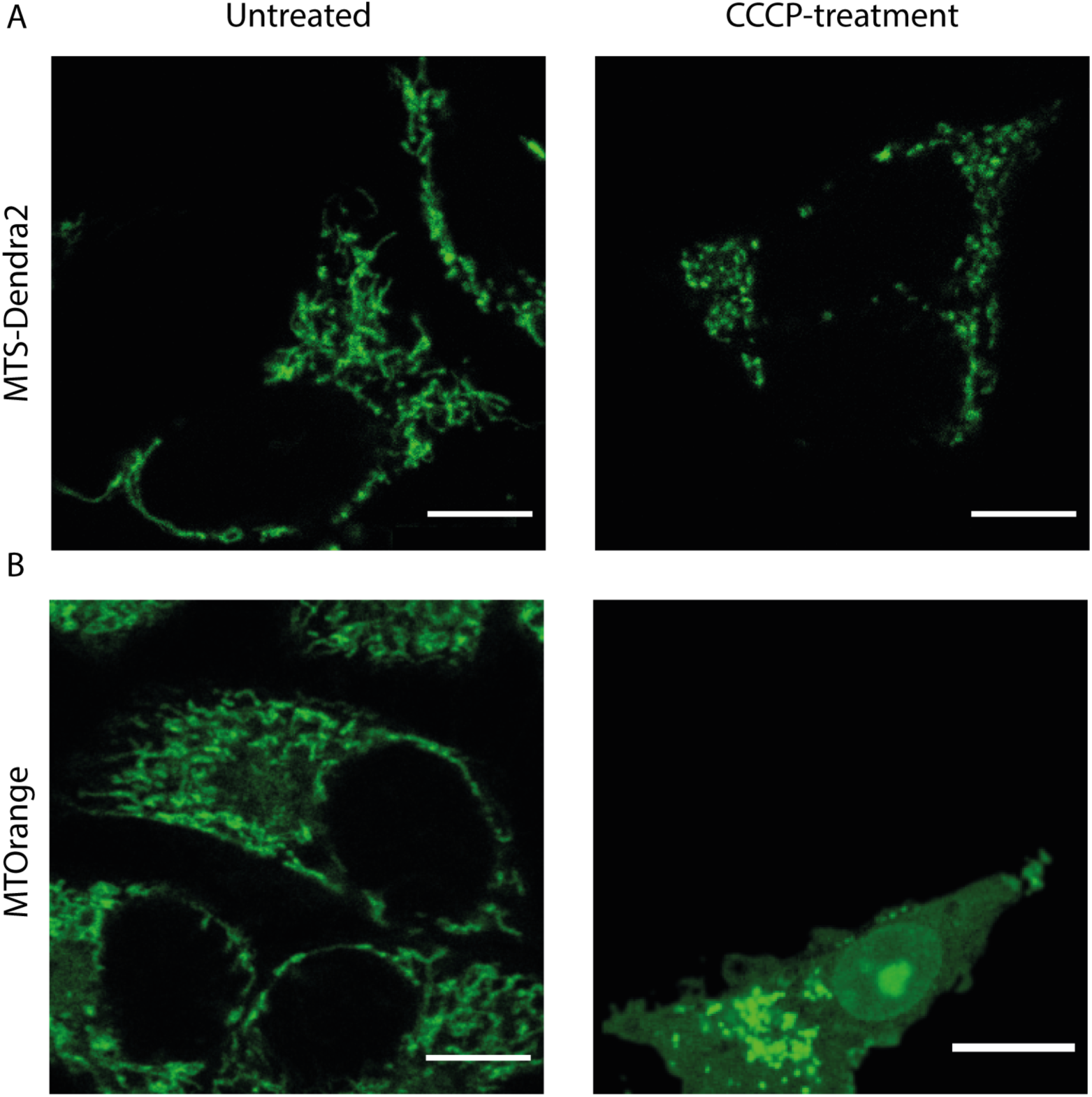
Mitochondria staining with the MTOrange and MTS-Dendra2 after CCCP treatment. Comparison between MTOrange and MTS-Dendra2 labeled cells under CCCP treatment. (A) The HEK 293 MTS-Dendra2 mitochondria (green). After 50 μm CCCP treatment mitochondrial network fragments, however, Dendra2 remains inside the mitochondria (B) The HeLa cells labeled with MTOrange (green). After 50 μm CCCP treatment, MTOrange partially exits the mitochondria, resulting in a low contrast of the fragmented mitochondria. Scale bar 10 μm.

## Supplementary videos

Video S1. pMC MTS-EmGFP expression vector injection into the HeLa cells.

The two-channel fluorescence time-series is represented by three windows in the video: the MTOrange labeled mitochondria fluorescence (green, left window), the MTS-EmGFP fluorescence (red, center window), and the merged fluorescence channel (right window). The time between the frames is 5 min.

Video S2. pMC MTS-Dendra2 expression vector injection into the HeLa cells.

The two-channel fluorescence time-series is represented by three windows in the video: the MTOrange labeled mitochondria fluorescence (green, left window), the MTS-Dendra2 fluorescence (red, center window), and the merged fluorescence channel (right window). The time between the frames is 5 min. The protein expression starts only after the completion of cell division about 4 h after the injection.

Video S3. MTS-EmGFP protein injection into the HeLa cells

The two-channel fluorescence time-series is represented by three windows in the video: the MTOrange labeled mitochondria fluorescence (green, left window), the MTS-EmGFP fluorescence (red, center window), and the merged fluorescence channel (right window). The time between the frames is 5 min. The MTS-EmGFP equilibrates between cytosol and nucleus within 1 h, however, the mitochondria continue to be seen as darker regions in the MTS-Em-GFP fluorescence image.

Video S4 MTS-SNAP-tag protein injection into the HeLa cells

The two-channel fluorescence time-series is represented by three windows in the video: the MTOrange labeled mitochondria fluorescence (green, left window), the MTS-SNAP-tag fluorescence (red, center window), and the merged fluorescence channel (right window). The time between the frames is 2 min. The MTS-SNAP-tag protein is imported into the mitochondria over the course of 1 h.

Video S5 MTS-SNAP-tag protein injection into the MTS-Dendra2 HEK 293 cell.

The two-channel fluorescence time-series is represented by three windows in the video: the MTS-Dendra2 labeled mitochondria fluorescence (green, left window), the MTS-SNAP-tag fluorescence (red, center window), and the merged fluorescence channel (right window). The time between the frames is 2 min. The MTS-SNAP-tag protein import into the mitochondria is seen after 1 h in the MTS-Dendra2 HEK 293 cell.

Video S6 SNAP-tag protein injection into the MTS-Dendra2 HEK 293 cell.

The two-channel fluorescence time-series is represented by three windows in the video: the MTS-Dendra2 labeled mitochondria fluorescence (green, left window), the SNAP-tag fluorescence (red, center window), and the merged fluorescence channel (right window). The time between the frames is 2 min. The SNAP-tag remains in the cytoplasm over the course of 1 h similar to the injection of the MTS-EmGFP into HeLa cell.

Video S7 MTS-SNAP-tag protein injection into the CCCP-treated MTS-Dendra2 HEK 293 cell

The two-channel fluorescence time-series is represented by three windows in the video: the MTS-Dendra2 labeled mitochondria fluorescence (green, left window), the MTS-SNAP-tag fluorescence (red, center window), and the merged fluorescence channel (right window). The time between the frames is 2 min. One noticeable feature in the cytoplasm is the appearance of bright dots of the MTS-SNAP-tag fluorescence. These peculiar dots are not co-localized with the mitochondria and are likely to be protein aggregates in lysosomes and proteasomes.

Video S8 MTS-SNAP-tag protein injection into the CHX-treated MTS-Dendra2 HEK 293 cell

The two-channel fluorescence time-series is represented by three windows in the video: the MTS-Dendra2 labeled mitochondria fluorescence (green, left window), the MTS-SNAP-tag fluorescence (red, center window), and the merged fluorescence channel (right window). The time between the frames is 2 min. The MTS-SNAP-tag protein import is clearly seen after 1 h.

Video S9 MTS-SNAP-tag protein injection into the PUR-treated MTS-Dendra2 HEK 293 cell

The two-channel fluorescence time-series is represented by three windows in the video: the MTS-Dendra2 labeled mitochondria fluorescence (green, left window), the MTS-SNAP-tag fluorescence (red, center window), and the merged fluorescence channel (right window). The time between the frames is 2 min. The mitochondria become noticeable in the protein channel at approximately 10 min and significantly brighter at 20 min, significantly faster compared to the import in the CHX-treated or untreated cells.

